# METTL3-mediated SOCS3 m^6^A Modification in Preserving CD4^+^ T Cell Homeostasis in HIV Infected Long-term Non-progressors

**DOI:** 10.1101/2025.11.13.688240

**Authors:** Jinming Su, Rongfeng Chen, Xing Tao, Sanqi An, Xiu Chen, Tongxue Qin, Yueqi Li, Jinmiao Li, Shanshan Chen, Yao Lin, Jie Liu, Jiemei Chu, Li Ye, Hao Liang, Junjun Jiang

## Abstract

The maintenance of normal CD4^+^ T cell levels in human immunodeficiency virus (HIV) infected long-term non-progressors (LTNPs) remains elusive. N6-methyladenosine (m^6^A) regulates RNA metabolism and immune function, but its role in LTNP pathogenesis is unelucidated. This study integrated in vivo analyses of peripheral blood mononuclear cells (PBMCs) from LTNPs, typical progressors (TPs), ART-treated patients, and healthy controls with in vitro experiments using the MT2-HIV-1_ⅢB_ model to explored the m^6^A regulatory role in LTNPs. Methylated RNA immunoprecipitation sequencing (MeRIP-seq) identified uniquely elevated m^6^A modification abundance and METTL3 expression in LTNPs vs. TPs. Differential m^6^A and mRNA analysis bighlighted enrichment of innate immunity/inflammation pathways, particularly the SOCS/IL/JAK axis. Further experiments confirmed that METTL3 mediates m^6^A modification of SOCS3, which suppresses JAK2/STAT4 phosphorylation to modulate IL-12/IFN-γ/IL-4 expression. These findings uncover a novel m^6^A-METTL3-SOCS3 regulatory axis underlying HIV long-term non-progression, explaining preserved CD4^+^ T cell homeostasis in LTNPs.

**IMPOTANCE**

LTNPs represent a unique subset of HIV-infected individuals who naturally maintain normal CD4⁺ T cell levels and slow disease progression, offering a valuable model to dissect host protective mechanisms against HIV. m⁶A modification has emerged as a pivotal regulator roles in HIV replication and T cell activation, yet its contribution to LTNP biology has remained unclear. This novel mechanism provides a potential explanation for the maintenance of normal CD4^+^ T cell levels in LTNPs. Our findings shed light on the molecular basis of HIV-1 long-term non-progression and offered crucial insights for developing functional cure for AIDS and new host immune strategies.

---

Human immunodeficiency virus (HIV) infection involves abnormal activation of the immune system, leading to excessive inflammation, gradual depletion of CD4^+^ T cells, and culminating in acquired immune deficiency syndrome (AIDS). Although the advent of antiretroviral therapy (ART)(1) has transformed AIDS into an incurable chronic disease, abnormal immune activation and inflammation persist, thereby influencing HIV disease progressions(2). A distinct subgroup of HIV-infected individuals, known as HIV-infected long-term non-progressors (LTNPs), accounting for approximately 5% of the HIV-infected population, maintain normal CD4^+^ T cell counts (>500 cells/uL) for over a decade without receiving ART(3). This cohort represents an essential model for investigating immune approaches and achieving a functional cure for AIDS.

The emergence of epitranscriptomics has provided new insights into AIDS therapeutics. N6-methyladenosine (m^6^A), the most abundant mRNA modification in eukaryotes(4), exhibits a dual role in viral immunity. It facilitates viral evasion while also stimulating host immune responses(5, 6). Studies have confirmed that HIV-1 genome undergoes m^6^A modification, a process that enhances viral replication and nuclear export(7). Methyltransferase-like 3 (METTL3), a core component of the m^6^A methyltransferase complex, modulates T cell proliferation/differentiation and immune responses(8, 9). Notably, Simona et al. demonstrated that METTL3 activators enhance viral particle production in HIV-1 provirus-carrying cells(10). Despite these advances, the intricacies of m^6^A-HIV replication, particularly in LTNPs, lacks clarity.

LTNPs exhibits a unique immunological profile featuring the preservation of normal CD4^+^ T cell counts(11). These CD4^+^ T cells encompass both T helper lymphocytes (Th) and regulatory T cells (Treg), each with distinct cytokine profiles and functions. Polarization of Th1/Th2 cell subsets has been shown to impact HIV disease progression(12, 13). Critically, m^6^A epigenetic modification regulates cell differentiation, and METTL3 has been implicated in maintaining T-cell balance. Mouse studies involving METTL3 knockouts models highlight its role in the peripheral naive CD4^+^ T cell proliferation and differentiation, suggesting its involvement in maintaining T cell homeostasis(14). The potential link between LTNPs’ immune balance and METTL3-mediated m^6^A modification of CD4^+^ T cells remains underexplored. This study innovatively explores the immune protective mechanism maintaining normal CD4^+^ T cell levels in LTNPs, from the perspective of m^6^A epigenetic regulation. By elucidating the role of m^6^A modification in mitigating HIV-induced inflammation and immune damage, the aim is to uncover novel therapeutic avenues for AIDS treatment.

## RESULTS

### Basic characteristics of study participants

A total of 12 subjects, comprising 3 LTNPs without receiving antiretroviral therapy (non-ART), 3 typical progressors (TPs) with non-ART, 3 ARTs, and 3 healthy controls (HCs), were recruited for Methylated RNA immunoprecipitation sequencing (MeRIP-seq). ANOVA revealed no statistically significant differences in age (*P*=0.487), gender (*P*=0.992), or ethnicity (*P*=0.802) across the four groups (Supplement Table S1).

### The m^6^A epitranscriptomic landscape

MeRIP-seq analysis of cleaned data revealed that the LTNPs group had a comparable number of m^6^A peaks to ARTs and HCs groups, and more than the TPs group. Furthermore, the distribution of m^6^A peaks in the 3’UTR region was higher in the LTNPs group than in the TPs group(Fig. 1A). Gene Ontology (GO) and Kyoto Encyclopedia of Genes and Genomes (KEGG) analyses of differential m^6^A peaks between LTNPs and TPs indicated their involvement in regulating type I interferon production, defense response to viruses, and the RIG-I-like receptor signaling pathway (Fig. 1B). RNA-seq data identified 1498 differentially expressed genes (DEGs) in the LTNPs versus TPs (Fig. 1C). Notably, these DEGs mainly focused on the regulation of T cell activation, immune effector processes, and antigen processing and presentation (Fig. 1D). Veen plots illustrated 24 overlapping genes between DEGs and differentially expressed m^6^A genes (DEm^6^A) in the LTNPs versus TPs (Fig. 1E). Linear correlation analysis demonstrated a negative association (*P*=0.030) between the transcriptome level of these overlapping genes and their m^6^A modification (Fig. 1F). Analyses of DEm^6^A and DEGs between LTNPs and the HCs/ARTs groups are shown in Supplement Fig. S1.

**FIG 1.**
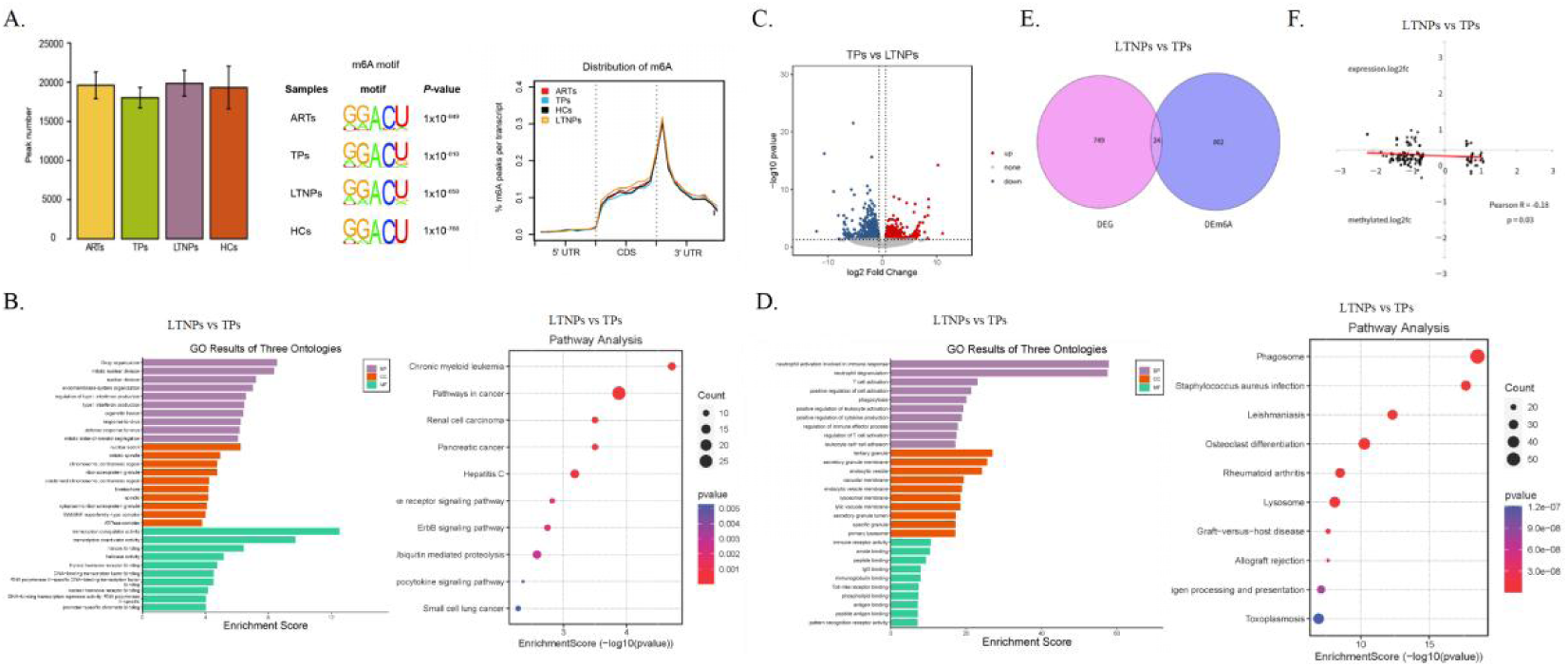
Map of m^6^A profiling in ARTs, TPs, LTNPs, and HCs groups. (A) On average, 19,615, 18,029, 19,870, and 19,324 m^6^A peaks were identified in the ARTs, TPs, LTNPs, and HCs group, respectively. The m^6^A motif was mainly enriched in RRACH consensus sequence across all four groups (R= A/G, H = A/C/U). The m^6^A peaks in each group of study populations are mainly distributed at the start of the 3’UTR region of the genome structure (5’UTR, CDS coding region, and 3’UTR), but the LTNPs group is significantly higher compared to the TPs group in the 3’UTR region. (B) GO and KEGG analyses of differential m^6^A peaks between LTNPs and TPs. (C) Volcano plots identified 1498 DEGs in the LTNPs versus TPs. (D) GO and KEGG analyses of DEGs between LTNPs and TPs. (E) Veen plots illustrated 24 overlapping genes between DEGs and differentially expressed m^6^A genes (DEm^6^A) in the LTNPs versus TPs. **F.** Linear correlation analysis demonstrated a negative association between the transcriptome level of these overlapping genes and their m^6^A modification (*P*=0.030). Note: Red represents upregulated mRNA, and blue represents downregulated mRNA in Volcano plots. In the bubble diagram, the size of bubble represents the number of genes in the target gene set that belong to a specific KEGG pathway, with larger bubbles indicating a higher number of genes.

### Relationship of HIV-1 and METTL3

Differential analysis of m^6^A regulatory factors demonstrated significant mRNA expression variations across the four groups, as depicted in the heatmap (*P*<0.05). Pairwise comparisons indicated upregulation of HNRNPD (*P*=0.002), TRA2A (*P*=0.006), METTL3 (*P*=0.007), ZCCHC4 (*P*=0.003), and RBM15 (*P*=0.031) in LTNPs compared to TPs (Fig. 2A). Population-level validation confirmed significantly higher METTL3 mRNA expression in LTNPs versus TPs (*P*=0.013), consistent with MeRIP-seq findings (Fig. 2B). mRNA expression of other m^6^A regulatory factors between LTNPs and TPs groups is presented in Supplement Fig. 2. To investigate the influence of HIV on METTL3 (Fig. 2C), both mRNA and protein levels of METTL3 were significantly reduced in the HIV-1 group compared to controls (*P*<0.05). Stable cell lines with METTL3 knockdown (sh-METTL3) and controls (sh-Vector) were successfully constructed (Fig. 2D) and validated by WB (*P*<0.001), the knockdown rate is 89%, indicating successful METTL3-knockdown. Upon infecting with the HIV-1_IIIB_ strain, HIV-1 p24 levels were significantly lower in the sh-METTL3 group compared to sh-Vector group at 12 hours (*P*=0.040), 24 hours (*P*<0.001), and 48 hours (*P*=0.004) post-infection (Fig. 2E).

**FIG 2.**
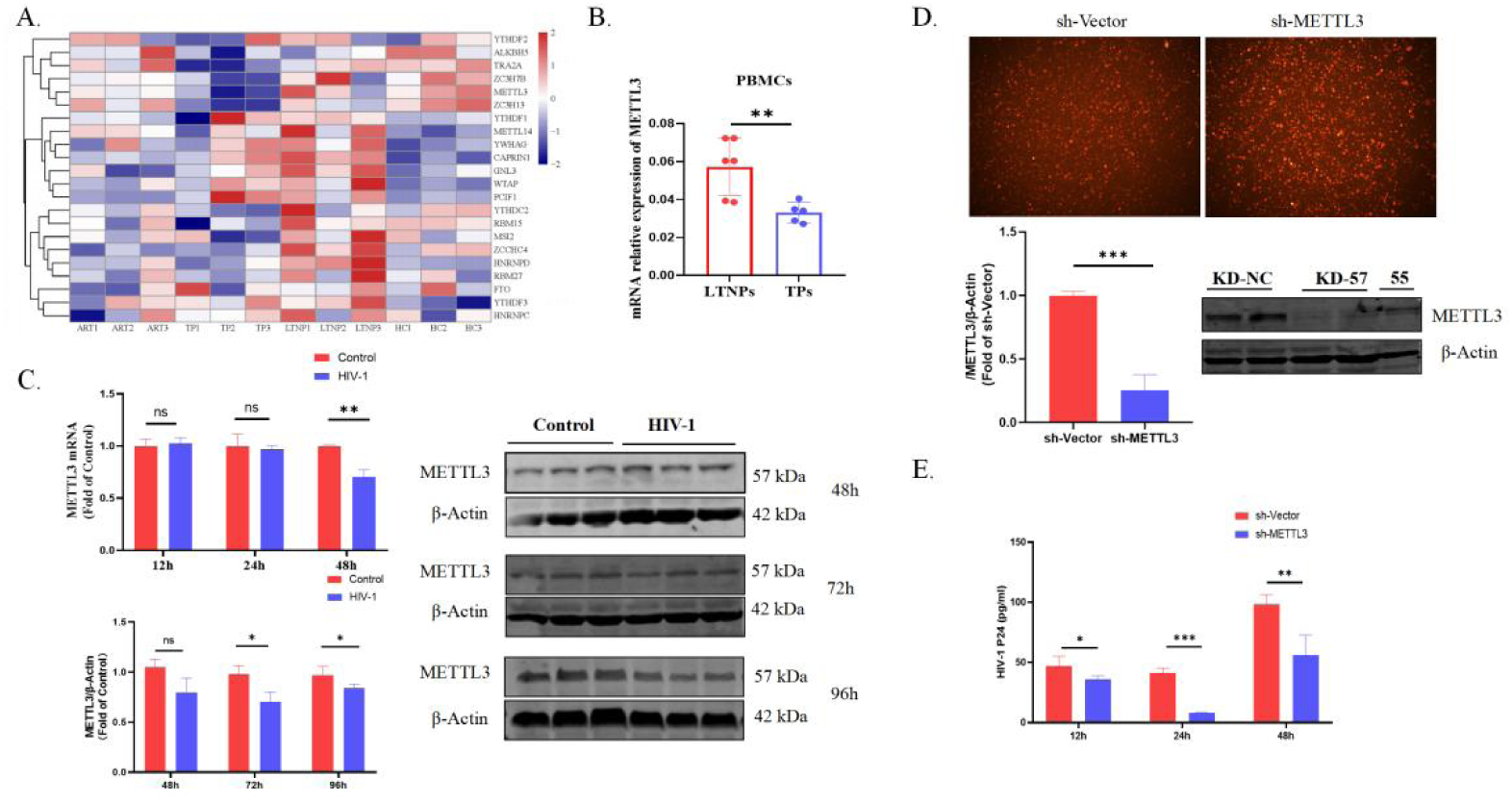
Relationship of HIV-1 and METTL3. (A) Pheatmap analysis demonstrated significant variations in m^6^A regulators mRNA levels across the four groups, including HNRNPD, TRA2A, METTL3, and so on (*P*>0.05). (B) Population-level validation confirmed that METTL3 mRNA expression was significantly higher in LTNPs than in TPs (*P*=0.013). (C) The METTL3 mRNA level at 48h was lower in HIV-1 infected group than in control group (*P*=0.002). There were no significant differences between the two groups at 12h and 24h (*P*=0.619, *P*=0.680). The METTL3 protein level at 72h and 96h were lower in HIV-1 infected group than in control group (*P*=0.013, *P*=0.041), There were no significant differences between the two groups at 24h and 48h (*P*=0.794, *P*=0.183). (D) Fluorescence images showed that lentiviruses for knockdown of METTL3 (sh-METTL3) and their negative control (sh-Vector) were transfected into MT2 cells. The METTL3 protein levels were lower in the sh-METTL3 group compared with sh-Vector.group (*P*<0.001). (E) Expression of HIV-1 p24 was lower in sh-METTL3 groups than in sh-Vector group at 12h, 24h, and 48h post-infection. (*P*=0.040, *P*<0.001, *P*=0.004)

### To verify the effect of METTL3 on SOCS3 expression

In-depth analysis uncovered differentially expressed inflammatory regulatory genes between LTNPs and TPs (*P*<0.05), with specific enrichment in the SOCS/IL/JAK pathway (Fig. 3A). Population-level validation confirmed higher SOCS3 expression in LTNPs compared to TPs (*P*=0.044) (Fig. 3B). Given SOCS3’s significance in HIV and T cell regulation(15, 16), The online database SRMAP was utilized to predict potential m^6^A modification sites on SOCS3 (Fig. 3C). Integrative Genomics Viewer (IGV) analysis revealed differential m^6^A peaks in IP productions between LTNPs and TPs (Fig. 3D). Our analysis, utilizing single-base elongation-and ligation-base qPCR amplification method (SELECT qPCR), verified a significant distinction in SOCS3-specific m^6^A sites between sh-METTL3 and sh-Vector (*P*=0.032), indicating that METTL3 target SOCS3 m^6^A sites in MT2 cells (Fig. 3E). Both qPCR and Western blot (WB) validations demonstrated downregulated SOCS3 mRNA and protein expressions in sh-METTL3-Control group versus sh-Vector-Control group under both uninfected and infected conditions (both *P*<0.001). These findings suggest that METTL3 promotes SOCS3 expression (Fig. 3F).

**FIG 3.**
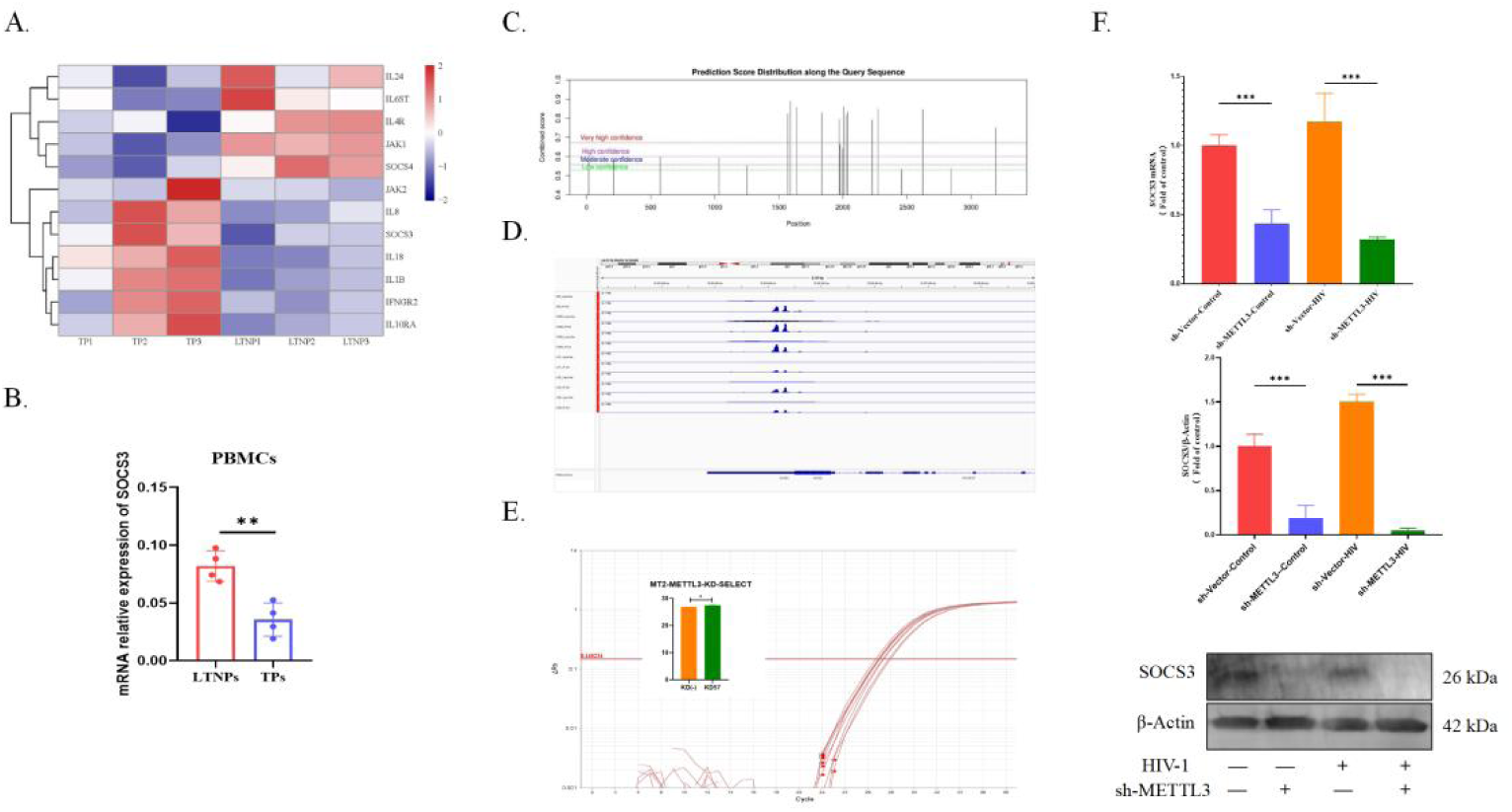
Effect of METTL3 on the expression of SOCS3. (A) Heatmap analysis depicted differentially expressed inflammatory-regulatory genes are enriched in SOCS/IL/JAK/STAT pathway between LTNPs and TPs groups. (B) SOCS3 mRNA expression was higher in LTNPs compared to TPs at population-level (*P*=0.044). (C) The online database SRMAP predict potential m^6^A modification sites on SOCS3, with the high credibility at the 1589, 1640, and 2006 positions. (D) Prediction of m^6^A sites in SOCS3 among LTNPs using IGV. (E) The SELECT qPCR experiment verified a significant distinction in SOCS3-specific m^6^A sites between sh-METTL3 and sh-Vector (*P*=0.032). (F) Both qPCR and WB validations demonstrated downregulated SOCS3 mRNA and protein expressions in sh-METTL3-Control group versus sh-Vector-Control groupunder both uninfected and infected conditions (both *P*<0.001). Note: - indicates HIV-1 un-infection or empty knockdown, + indicates HIV-1 infection or METTL3 knockdown

### The effect of METTL3 and SOCS3 on the regulation of IL-12, IFN-γ, and IL-4

In Fig. 4A, ELISA analysis revealed that METTL3 knockdown significantly upregulated IL-12 protein expression (*P*<0.001) and downregulated IL-4 expression (*P*=0.007) compared to the control. After HIV-1 infection, these trends persisted, with METTL3 knockdown maintaining higher IL-12 levels (*P*<0.001) and lower IL-4 levels (*P*=0.002). No significant difference in IFN-γ protein demonstrated was observed between the two groups (*P*>0.05). Subsequently, stable SOSC3-overexpression (SOCS3-OE(+)) and negative control (SOCS3-OE(-)) MT2 cell lines were established (Fig. 4B), and validated by WB (*P*<0.001). Furthermore, HIV-1 p24 levels were significantly higher in the SOCS3-OE(+) group than in the SOCS3-OE(-) group (*P*=0.021) (Fig. 4C). ELISA analysis showed that in uninfected cells, SOSC3 overexpression significantly reduced IL-12 and IFN-γ expression compared to the control (both *P*<0.001). Upon HIV-1 infection, SOCS3-OE(+) group exhibited significantly decreased IL-12 and IFN-γ levels (*P=*0.042, *P*=0.001), along with increased IL-4 expression (*P*=0.012), relative to the SOCS3-OE(-) group (Fig. 4D). These findings suggest that METTL3-mediated SOCS3 regulates IL-12/IL-4 signaling during HIV infection.

**FIG 4.**
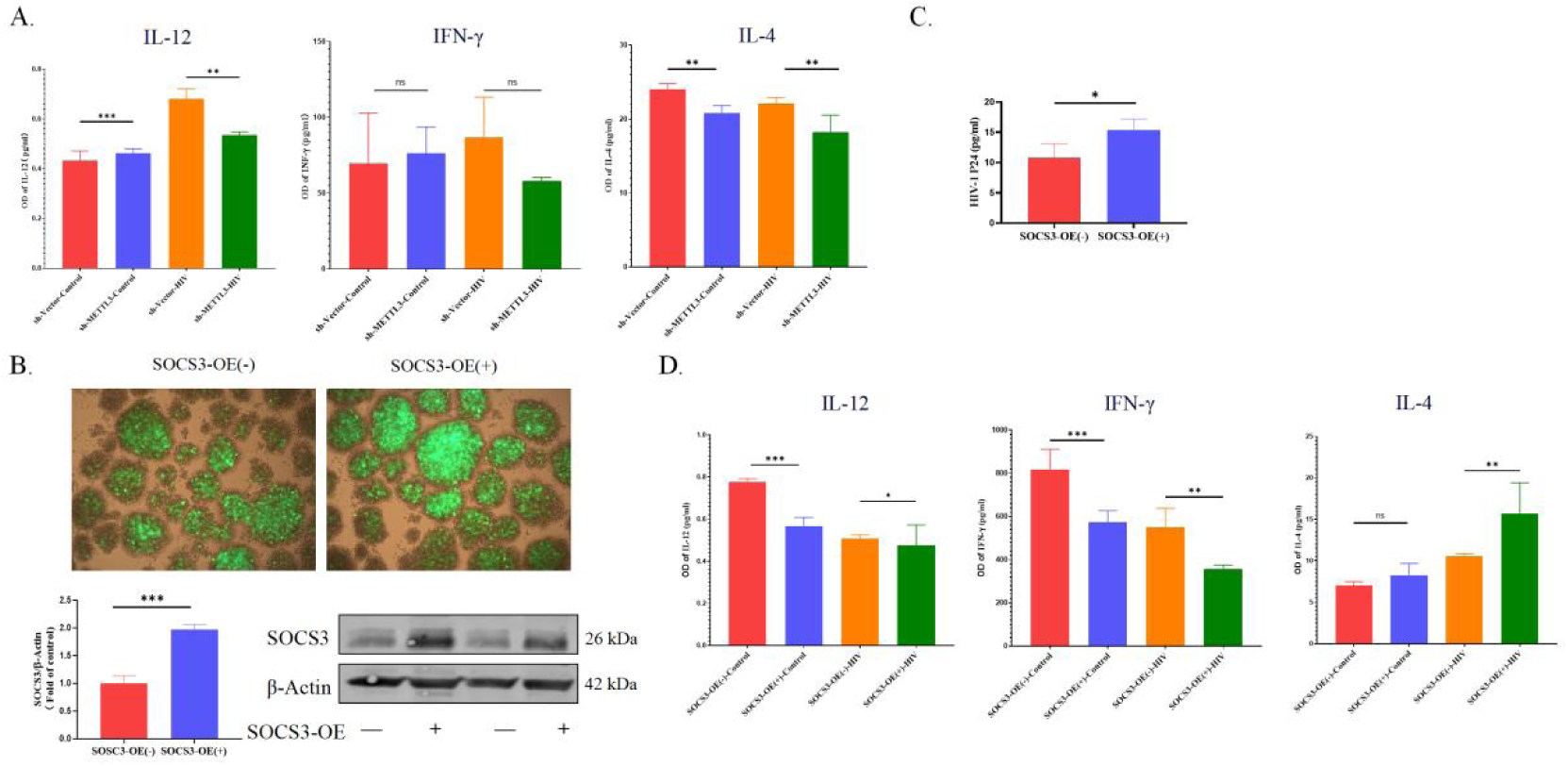
The Effect of METTL3 and SOCS3 on IL-12, IFN-γ, and IL-4. (A) METTL3 knockdown significantly upregulated IL-12 protein (*P*<0.001, *P*<0.001) and downregulated IL-4 expression (*P*=0.007, *P*=0.002) under both uninfected and infected conditions. (B) Fluorescence images showed that lentiviruses for overexpression SOSC3 (SOCS3-OE (+)) and their negative control (SOCS3-OE (-)) lentivirus were transfected into MT2 cells. The SOSC3 protein levels were higher in the SOCS3-OE (+) group compared with (SOCS3-OE (-) group (*P*<0.001). (C) HIV-1 p24 levels were significantly higher in the SOCS3-OE(+) group than in the SOCS3-OE(-) group (*P*=0.021). (D) In both uninfected and infected cells, SOCS3 overexpression significantly reduced IL-12 protein (*P*<0.001, *P*=0.042) and IFN-γ protein (*P*<0.001, *P*=0.001) compared to the control group, while increased IL-4 protein levels were observed only in infected cells (*P*=0.012). Note: - indicates HIV-1 un-infection or empty overexpression, + indicates HIV-1 infection or SOCS3 overexpression

### Exploring METTL3’s regulation of the inflammatory response via the JAK2/STAT4 pathway

SOCS3 is a key regulatory factor in the JAK/STAT pathway, through which IL-12 and IL-4 also exert their inflammatory effects. Further research was conducted to investigate whether METTL3-mediated m^6^A modification of SOCS3 influences these inflammatory factors via the JAK/STAT pathway. Under both HIV-uninfected and infected conditions, JAK2 mRNA expression was higher in the sh-METTL3 group than in the sh-Vector group (*P*=0.045, *P*=0.015). STAT4 mRNA levels exhibited the same trend (both *P<*0.001), indicating that METTL3 can inhibit JAK2 and STAT4 mRNA expression (Fig. 5A). Additionally, JAK2 phosphorylation, which is crucial for its activation, subsequently mediates STAT4 autophosphorylation. Comparative analysis revealed no significant difference in JAK2 protein levels between METTL3-knockdown and control groups (*P*=0.970). However, phosphorylated STAT4 protein levels were higher in METTL3-knockdown uninfected cells (*P*=0.005). After HIV-1 infection, METTL3 knockdown significantly elevated phosphorylated JAK2 and STAT4 protein levels compared to the control group (*P*<0.001, *P*<0.001), suggesting that METTL3 activates JAK2 and subsequently promotes phosphorylated STAT4 protein expression in HIV-infected MT2 cells (Fig. 5B).

**FIG 5.**
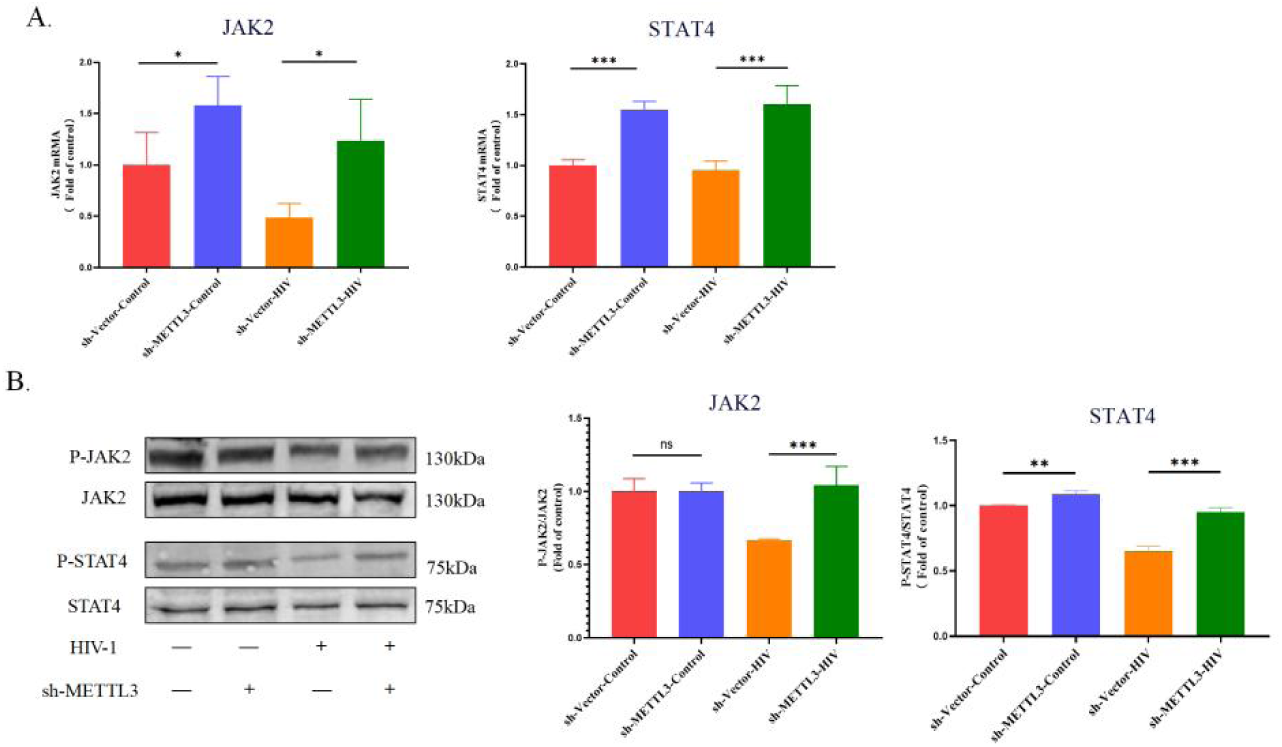
Investigating the modulatory role of METTL3 on the inflammatory response via the JAK2/STAT4 axis. (A) Under both HIV-uninfected and infected conditions, JAK2 mRNA expression was higher in the sh-METTL3 group than in the sh-Vector group (P=0.045, P=0.015). STAT4 mRNA levels exhibited the same trend (both P<0.001). (B) METTL3 knockdown significantly elevated phosphorylated JAK2 (*P*<0.001) and STAT4 (*P*<0.001) protein levels compared to the control group after HIV-1 infection.

## DISCUSSION

LTNPs exhibit distinct genetic and immunological characteristics that offer promising insights for developing potential immune pathways, therapeutic targets, and vaccines strategies against HIV(17, 18). While m^6^A modification is intricately involved in HIV replication(7), its specific role in LTNPs remains poorly understood. MeRIP-seq profiling uncovered a unique m^6^A epitranscriptomic landscape in LTNPs.

Furthermore, functional enrichment analyses highlighted DEm^6^A and DEGs involved in T cell activation and immune effector processes in LTNPs versus TPs. Notably, LTNPs showed a balanced Th1/Th2 T-cell profile, consistent with established literature(13, 19). Differentially expression of key inflammatory regulatory (e.g., SOCS3, IFNGR2, IL-4R, JAK1/2) between these groups were also observed. Importantly, mRNA-m^6^A methylation analysis elevated METTL3 expression in LTNPs, a finding validated in a separate cohort. Given the documented role of METTL3 in T-cell homeostasis and differentiation(20-22), These data suggest that the m^6^A modifications, mediated by METTL3, contribute to unraveling the distinctive immunoregulatory phenotype of LTNPs.

The balance between Th1 and Th2 cytokines is critical for immune regulation. IL-12, a key Th1 cytokine, drives naïve T cell differentiation and possesses dual proinflammatory and immunomodulatory activities(23), with overexpression linked to autoimmunity. Conversely, IL-4, a hallmark of Th2 responses(24), promotes CD4^+^ T cells activation and Th2 differentiation, thereby maintaining immune homeostasis and restraining excessive inflammation(25, 26). This study demonstrated that METTL3 inhibited IL-12 expression while upregulating IL-4 levels (*P*<0.05), suggesting its role in mitigating inflammatory responses. This shift in the cytokine milieu may contribute to immune protection in LTNPs, warranting further exploration of METTL3-mediated mechanisms.

As a key negative regulator of the JAK/STAT pathway(27), SOCS3 influences T-cell balance and HIV replication(28, 29). SELECT qPCR results confirmed that METTL3 mediated m^6^A modification of SOCS3(30), and its knockdown significantly decreased SOCS3 levels (*P*<0.001), validating METTL3’s promotive effect(31). To delineate the functional impact, we found that SOCS3 overexpression suppressed IL-12 expression (*P*<0.05), consistent with its known blockade of the IL-12/STAT4 pathway(32). Since IL-12 stimulated IFN-γ(33), which could exacerbate inflammation via TNF-ɑ and IL-6(34), SOCS3 concurrently attenuated IFN-γ expression (*P*<0.05). Conversely, overexpression-SOCS3 enhanced IL-4 levels post-infection (*P*=0.012). This establishes a direct link between epitranscriptomic regulation and cytokine signaling, suggesting that the METTL3-SOCS3 axis helps mitigate excessive immune activation in LTNPs.

Having established the role of METTL3 in regulating the SOCS3-cytokine axis, subsequent exploration focused on its impact on downstream JAK/STAT signaling. Although METTL3 has been reported to activate JAK2/STAT3 and then leading to skin lesions(35), in HIV-1 infection, it was found to suppress JAK2 expression and phosphorylation (*P*<0.001). Since IL-12 and IFN-γ activate JAK/STAT signaling(36-38) and given the specific role of STAT4 in mediating inflammatory responses(39, 40), METTL3 knockdown was found to augmented STAT4 phosphorylation (*P*<0.001). These data indicated that METTL3 exerts an inhibitory effect on JAK2/STAT4 pathway. Integrating these findings, this study proposes a novel mechanism: METTL3-mediated m^6^A modification upregulates SOCS3 expression, which consequently shifts the cytokine balance from IL-12/IFN-γ toward IL-4, potentially explaining the mitigated immune activation observed in LTNPs.

This study has several limitations. Technically, site-specific mutagenesis of SOCS3 were hampered by difficulties in maintaining MT2 cell growth and achieving high-efficiency mutation, resulting in insufficient specificity. Clinically, the current standard of immediate treatment initiation limited to validate the METTL3-SOCS3 regulatory axis with MeRIP-RNA assays in a larger cohort.

MeRIP-seq analysis identifies a distinct m^6^A epitranscriptomic landscape in CD4^+^T cells of LTNPs, providing crucial insights into the mechanism of m^6^A in HIV pathogenesis. Furthermore, a novel immunoregulatory pathway is delineated in which METTL3-mediated m^6^A modification of SOCS3 modulates the IL-12/STAT4 axis, thereby contributing to preserved CD4^+^ T-cell homeostasis and delayed disease progression in LTNPs. These discoveries not only help to elucidate the molecular mechanism of HIV long-term non-progression but also offer potential insights for developing therapeutic strategies aimed at achieving a functional cure for AIDS.

## MATERIALS AND METHODS

### Study participants

Participants aged 18 years or older, residing in Guangxi, confirmed HIV positive, and who had not progressed to AIDS were recruited. All study participants were divided into four groups based on different definitions, namely LTNPs, TPs, ARTs, and HCs. LTNPs were defined as HIV-positive for ≥10 years with stable CD4^+^ T cell counts (>500 cells/μL), without receiving ART. TPs were HIV-positive for 3-9 years with declining CD4^+^ T cell counts to <500 cells/μL without receiving ART. ARTs comprised individuals on ART for >1 year with viral load <1000 copies/mL. Additionally, HIV-negative individuals constituted the HCs group. Study participants were recruited in Guangxi between January and March 2021 based on matching of age, gender, and ethnicity. All subjects are approved by the Ethics Committee of Guangxi Medical University [review 20210092], following the principle of informed consent and signing an informed consent form.

### Materials

MT2 cells were derived from the transformation induced by human T-lymphotropic virus type I. HIV-1_ⅢB_ stain was owned by the Biosafety Level 3 Laboratory of Guangxi Medical University. Primary antibodies included Anti-β-Actin antibody, METTL3 (E3F2A) Rabbit monoclonal antibody (mAb), SOCS3 (D6E1T) Rabbit mAb, Janus kinase 2 (JAK2) (D2E12) Rabbit mAb, Phospho-JAK2 (Tyr1007/1008) (C80C3) Rabbit mAb, Signal transducer and activator of transcription 4 (STAT4) (C46B10) Rabbit mAb, Phospho-STAT4 (Tyr693) (D2E4) Rabbit mAb. Secondary antibodies included Donkey-anti mouse 680RD and Donkey-anti rabbit 800CW.

### Isolation of peripheral blood mononuclear cells (PBMCs)

10 mL of peripheral blood was collected and centrifuged at 800 g for 5 minutes at 20℃. The lower layer of blood cells was used for PBMC isolation. Added 20 mL of Ficoll to a tube, layered diluted blood cells (1:1 with PBS) and centrifuged at 1200 × g for 10 minutes at 20℃. The grayish-white layer of PBMCs was collected after washing and centrifuging. Then, erythrocyte lysate was added, let stand for 10 min, and terminated with PBS.

## Magnetic bead sorting of CD4^+^ T lymphocytes

We resuspended 1×10^7^ cells PBMCs in a mixture of 80 μL of running buffer and 20 μL of CD4 microbeads. These suspension was incubated on ice for 15 minutes, ensuring the absence of bubbles. Following incubation, the cells were washed, centrifuged, and resuspended in 0.5 mL of running buffer. The sorting column (LS) was installed on a magnetic rack, with a 15 mL centrifuge tube (negative collection) underneath. The LS was rinsed with 3 mL of running buffer. The suspended PBMCs were added to the LS, and the LS was rinsed three times with 3 mL of running buffer each time. The LS was transferred to a new 15 mL tube (positive collection) containing 5 mL of running buffer. The plunger was immediately pushed into the column to obtain bead-labeled CD4^+^ T lymphocytes. The collected cells were resuspended and counted by centrifugation at 300 × g for 5minutes at 20 ℃. The purified CD^+^ T cells were stored in EP tubes for RNA extraction.

### Extraction of total RNA

We incorporated chloroform (1 : 5 dilution with Trizol) into the above EP tube, vortexed it for 15 seconds, and incubated at 20℃ for 10 minutes. Subsequently, the tube was centrifuged at 14,000 × g for 15 minutes at 4°C. The supernatant was transferred to a fresh tube, and isopropanol (1 : 2 dilution with Trizol) was added, followed by incubation and centrifugation. The RNA was then washed with 75% ethanol (1 : 1 dilution with Trizol) and dissolved in 20 μL of DEPC. RNA quality was measured using NanoDrop, with an A260/A280 > 1.8 deemed suitable for further use.

### MeRIP-seq

Following total RNA extraction, the enriched RNA samples were fragmented into short fragments (approximately 100 nt) using a fragmentation buffer. The fragmented RNA was divided into two portions, one served as the input (without immunoprecipitation), while the other was subjected to immunoprecipitation (IP) using an anti-m^6^A specific antibody. The enriched RNA was then reverse-transcripted into cDNA using random primers. Subsequently, the cDNA fragments were end-repaired and ligated to Illumina sequencing adapters. Finally, qualified library were constructed and sequenced on the Illumina NovaSeq^TM^ 6000. Raw sequencing data were processed with FASTQC for quality assessment, followed by the removal of adapters, duplicate, and low-quality sequences to generate clean data.

### m^6^A analysis

For m^6^A data analysis, peak calling, annotation, and motif analysis were performed by comparing the m^6^A-IP and input samples. Difference m^6^A peaks were identified using the thresholds FDR<0.05 and |log2FC|>1. Ultimately, by integrating RNA-seq data and conducting GO and KEGG enrichment analysis, the potential regulatory functions of m^6^A modification in gene expression and associated biological pathways were investigated.

### MT2 cells thawing and culture

We formulated a complete culture medium of PRMI 1640 supplemented with 10% FBS and 1% double antibiotics (comprising 100 U/mL penicillin and 100 μg/mL streptomycin). Subsequently, 1 mL of MT2 cells were added to 5 mL of this medium in a tube, thoroughly mixed, and centrifuged at 900 rpm/min for 5 minutes at 20℃. The supernatant was discarded, and the cell suspension was resuspended in 1 mL of PRMI 1640 before being transferred into a 25 T culture flask. The cells were cultured at 37°C and 5% CO^2^ until they reached a stable third generation, at which point they were prepared for staining or transfection experiments.

### HIV-1_IIIB_ infection and amplification in MT2 cells

5 mL of logarithmically proliferating MT2 cells (2×10^5^ cells/mL) were cultured in a 25 T flask with 1 mL of HIV-1_ⅢB_ supernatant at 37°C and 5% CO^2^. Daily monitoring encompassed medium hue, cellular status, and cytopathic effect (CPE). On day 3, 3 mL of fresh PRMI 1640 was substituted. Samples were then collected at various intervals or upon substantial cell mortality. The supernatant, containing the HIV-1_ⅢB,_ was centrifuged at 1500 rpm/min for 10 minutes at 4 °C, quantified for the tissue culture infective dose 50% (TCID50), and stored at -80°C.

### Construction of lentivirus transfection-MT2-HIV-1_ⅢB_ infected cell model

Logarithmically growing MT2 cells (5×10^5^ cells/mL) were inoculated in a 48-well plate containing PRMI 1640 devoid of FBS and supplemented with lentiviral infection enhancers. Lentivirus were introduced to MT2 cells according to cells density, lentiviral titer, and MOI (10, 30, 50), follow by incubation at 37℃ and 5% CO^2^ for 12-16 hours. Infection efficiency was assessed after 72 hourspost-inoculation. Upon achieving 80% infection efficiency, cells were maintained in 2 μg/mL Puromycin, with media exchanges every 3 days o eliminate non-transduced cells. Once 100% efficiency was confirmed, Puromycin concentration was reduced to maintenance level, and overexpression/knockdown was verified using qPCR and WB. Subsequently, cell lines and negative controls were incubated with HIV-1_ⅢB_ strain (p24 20 ng/10^6^ cells) under 37℃ and 5% CO^2^ conditions for 3 hours. The cells were then centrifuged at 2500 rpm/min for 5 minutes at 20°C, resuspended in FBS, and further cultured in medium for collecting samples at different time points for follow-up experiments.

### qPCR

Total RNA reversed transcribe cDNA using the TAKARA RR036A. The qPCR quantification was performed by TAKARA RRR820A, using the Primer Bank database (https://pga.mgh.harvard.edu/primerbank/) to design primers, as shown in Supplement Table S2. The reaction system and condition of reverse transcription and qPCR were performed according to the instructions.

### WB

We dispensed 10 mL of 10% resolving gel into a glass plate, sealed the surface with ddH2O, and allowed it to solidify for 20 minutes. Subsequently, the excess water was removed, and 4 mL of stacking gel was added onto the resolving gel. A 10-well comb was inserted, and the gel was allowed to solidify completely. The comb was then removed vertically, and the gel was filled with electrophoresis buffer. Each well was loaded with 3 μL of rainbow marker and 40-60 μg of protein samples. Electrophoresis was performed at 80-120 V for 1.5 hours, or until the indicator reached the bottom of the gel. Following electrophoresis, the proteins were transferred onto a PVDF membrane using a "sandwich" transfer assembly (sponge-filter paper-gel-membrane-filter paper-sponge) at 250 mA for 60 minutes. The PVDF membrane was washed with TBST and incubated with the primary antibody and secondary antibody. Finally, the PVDF membrane was scanned using an Odyssey CLX dual-color infrared laser imaging system, and the resulting images were analyzed using Image J software.

### ELISA for HIV-1 p24 antigen and host cytokines

We equilibrated the HIV-1 p24 ELISA kit at 20 ℃ for 1.5 hours prior to use. Cell supernatants were thawed and diluted 1000-fold to ensure thorough mixing. Experimental setup included control, duplicate, and blank wells. Subsequently, 25 μL/well of lysate (excluding blank wells), 100 μL/well of standards, and samples were added, followed by incubated with shaking at 37℃ for 1 hours. The plate was then washed 5 times, and 125 μL of HRP-avidin was added to each well, followed by another incubation with shaking at 37℃ for 1 hour. Termination solution (50 μL/well) was added, and the optical density (OD) was measured at 450 nm (with 630 nm as the reference wavelength) within 5 minutes. Lastly, A standard curve was plotted to calculate the HIV-1 p24 levels. Additionally, IL-12, IFN-γ, and IL-4 ELISAs were performed adhering strictly to the kit manufacturer’s instructions.

### SELECT qPCR

We used the Epi-SELECTTM m^6^A site identification kit to detect m^6^A modification. Designed Up/Down probes of m^6^A based on MeRIP Seq. The gene-specific primers were designed for qPCR to ensure that the initial RNA of the control and experimental groups was the same. RNA template pretreatment (10 pg to 5 ug), heated at 65 °C for 5 min, then placed on ice for 3 min to denature RNA. Synthesized the first-strand cDNA, and performed qPCR to quantify the m^6^A site using the 2^-ΔΔCT^ method. The annealing extension was continued for the target m^6^A site, and then single base extension ligation was performed and verified by qPCR using TaKaRa RR820A containing Select F/R primers. The SELECT qPCR primers were specifically designed to target m^6^A sites within target genes, which conserved RRACH sequence of these peaks (Supplement Table S3).

### Statistical analysis

The study utilized IBM SPSS 26.0 (SPSS Inc. Chicago, USA) for analysis, with results graphically presented using GraphPad Prism 9.5 software. The t-test and analysis of variance (ANOVA) were employed for comparison, all statistical tests were two-sided with a significance level of *P*<0.05.

## ACKNOWLEDGMENTS

We thank Dr. Shan Lu of the University of Massachusetts Medical School for kindly providing us with the MT2 cells. We also gratefully acknowledge the Center for Disease Prevention and Control of Liuzhou, Guigang, Qinzhou, Yulin, and Laibing for collecting and providing epidemiological data on study populations.

## AUTHORS CONTRIBUTIONS

Junjun Jiang, Hao Liang, and Li Ye conceived and designed the study│Jinming Su, Rongfeng Chen, and Xing Tao performed the overall experiments and wrote the original manuscript│Xiu Chen, Tongxue Qin, Jinmiao Li, and Shanshan Chen help to carry out the samples experiments and interpreted data│Sanqi An, Yueqi Li, and Yao Lin contributed to bio-informatics design and analysis│Jie Liu and Jiemei Chu supervised the study. All authors approved the final version of the manuscript to be published.

## DATA AVAILABILITY

The HIV patient datasets generated and/or analyzed during the current study are not publicly available for ethical and legal reasons. Still, they are available from the corresponding author on request.

## DECLARATIONS

### CONSENT FOR PUBLICATION

Not applicable.

## Supplemental Material

**Supplemental material Table S1** The characteristics of each group.

**Table S2** The main primer sequence of qPCR.

**Table S3** Primer sequences of SELCET qPCR

**FIG S1** Analyses of DEm^6^A and DEGs between LTNPs and HCs/ARTs groups. (A) GO and KEGG analyses of differential m^6^A peaks between LTNPs and HCs indicated involvement in histone modification, transcription coactivator activity, and Fc gamma R−mediated phagocytosis. For LTNPs vs. ARTs, differential m^6^A peaks were associated with negative regulation of phosphorylation, ubiquitin−like protein transferase activity, and apoptosis. (B) Volcano plots identified 1200 and 1006 DEGs in the LTNPs group compared to the HCs and ARTs, respectively. (C) GO and KEGG analyses showed that DEGs between LTNPs and HCs mainly focused on the T cell activation, immune receptor activity, antigen processing and presentation. For LTNPs vs. ARTs, DEGs were enriched in interferon−gamma−mediated signaling pathway, MHC protein complex, and antigen processing and presentation. (D) Veen plots illustrated 94 and 43 overlapping genes between DEGs and DEm^6^A in the LTNPs ompared to the HCs and ARTs, respectively. (E) Linear correlation analysis demonstrated no significant association between transcriptome levels and m^6^A modification in LTNPs versus HCs or ARTs (*P* >0.05).

**FIG S2** mRNA expression of m^6^A regulatory factors between LTNPs and TPs groups in population-level. YTDHF3 (*P*=0.039), YTHDC2(*P*=0.008), and ALKBH5(*P*=0.023) mRNA expression were significantly higher in LTNPs than in TPs In contrast, no significant differences in mRNA expression were observed for other m^6^A regulatory factors between the two groups (*P*>0.05), including METTL4, TRA2A, YTHDF1, and so on.

